# Natural selection on plasticity of thermal traits in a highly seasonal environment

**DOI:** 10.1101/191825

**Authors:** Leonardo D. Bacigalupe, Juan D. Gaitan-Espitia, Aura M. Barria, Avia Gonzalez-Mendez, Manuel Ruiz-Aravena, Mark Trinder, Barry Sinervo

**Affiliations:** Instituto de Ciencias Ambientales y Evolutivas, Facultad de Ciencias, Universidad Austral de Chile, Casilla 567, Valdivia, Chile; The Swire Institute of Marine Science and School of Biological Sciences, The University of Hong Kong, Hong Kong SAR, China; CSIRO Oceans and Atmosphere, GPO Box 1538, Hobart 7001, TAS, Australia; School of Natural Sciences, College of Sciences and Engineering, University of Tasmania, Hobart, Tasmania, Australia; MacArthur Green, 95 South Woodside Road, Glasgow, UK; Department of Ecology and Evolutionary Biology, University of California, Santa Cruz, CA, 95064, USA

**Keywords:** Amphibians, natural selection, physiological plasticity, acclimation, *Pleurodema thaul*, Atacama Desert

## Abstract

This preprint has been reviewed and recommended by Peer Community In Evolutionary Biology (http://dx.doi.org/10.24072/pci.evolbiol.100048). For ectothermic species with broad geographical distributions, latitudinal/altitudinal variation in environmental temperatures (averages and extremes) are expected to shape the evolution of physiological tolerances and the acclimation capacity (i.e., degree of phenotypic plasticity) of natural populations. This can create geographical gradients of selection in which environments with greater thermal variability (e.g., seasonality) tend to favour individuals that maximize performance across a broader range of temperatures compared to more stable environments. Although thermal acclimation capacity plays a fundamental role in this context, it is unknown whether natural selection targets this trait in natural populations. Here we addressed such an important gap in our knowledge by measuring survival, through mark recapture integrated into an information-theoretic approach, as a function of the plasticity of critical thermal limits for activity, behavioural thermal preference and the thermal sensitivity of metabolism in the northernmost population of the four-eyed frog *Pleurodema thaul*. Overall, our results indicate that thermal acclimation in this population is not being targeted by directional selection, although there might be signals of selection on individual traits. According to the most supported models, survival decreased in individuals with less tolerance to cold when cold-acclimated (probably because daily low extremes are frequent during the cooler periods of the year) and increased with body size. However, in both cases, the directional selection estimates were non-significant.

## Introduction

It is well known that environmental temperature (T_a_) is the abiotic factor with major incidence in the evolution, ecology and physiology of most of the biodiversity in the planet (Angilletta 2009 and references therein). The effects of T_a_ are particularly relevant for ectotherms as their body temperature depends on T_a_ and therefore any change in T_a_ affects their fitness and performance (e.g. behaviour, growth, reproduction, metabolism). This relationship between performance and temperature has been described by a thermal performance curve (TPC) (Huey & Berrigan 2001; Angilletta 2009) which has often been used to describe the thermal ecology and evolution of ectotherms (Gilchrist 1995; Huey & Kingsolver 1989), their phenotypic plasticity (Schulte et al. 2011), and to predict their responses to climate change (Clusella-Trullas et al. 2011; Sinclair et al. 2016). The TPC is best captured by three parameters: a minimum critical temperature (CT_Min_), which represents T_a_ below which performance is minimum; a maximum critical temperature (CT_Max_), which represents T_a_ above which performance is also minimum and an optimum temperature (T_Opt_), which represents T_a_ at which performance is maximum. Most of these parameters can exhibit geographic variation depending on the particular environmental context (e.g., local climate) and genetic background of populations (Gilchrist 1996; Kingsolver et al. 2004; Latimer et al. 2011). Furthermore, this geographic variation has the potential to create gradients of selection for TPCs across the species distribution (Kingsolver & Gomulkiewicz 2003) shaping thermal sensitivities, tolerances and thermal acclimation capacities (i.e., thermal plasticity) of local populations (Seebacher et al. 2012; Gaitan-Espitia et al. 2014).

Different climate-related hypotheses have been proposed to explain how physiological tolerances, capacities and their plasticity affect the distributional ranges of species (Bozinovic et al. 2011). One of them, the climate variability hypothesis (CVH), offers a powerful conceptual framework to explore the interactions between environmental variability and physiological performance of ectotherms (e.g., Gaitan-Espitia et al. 2013; 2014). The CVH predicts that organisms inhabiting more variable environments should have broader ranges of environmental tolerance and/or greater physiological plasticity that enable them to cope with the fluctuating environmental conditions (e.g., seasonality) (Ghalambor et al. 2006; Gaitan-Espitia et al. 2017). In agreement with this hypothesis, other theoretical models have explored the evolutionary mechanisms underlying local thermal adaptation across heterogeneous environments (e.g., Generalist-Specialist models). For instance, the model developed by Lynch and Gabriel (1987), predicts that temporal environmental heterogeneity selects for more broadly adapted individuals, whereas in more constant environments the model developed by Gilchrist (1995), predicts that selection should favor thermal specialists with narrow performance breadth. The mechanistic understanding of these conceptual frameworks has improved with recent studies showing how in thermally variable environments directional selection acts on TPC’s parameters favoring organisms that maximize performance across a broader range of temperatures (Logan et al. 2014) despite the ability of ectotherms to thermoregulate behaviorally (Buckley et al. 2015). Notwithstanding this progress, whether natural selection targets thermal acclimation capacity (i.e., plasticity) itself in natural populations remains unknown.

In addition to increasing mean temperatures, it is known that climate change is changing the frequency and intensity of extreme temperatures and events (Rahmstorf & Coumou 2011; Wang & Dillon 2014; Vazquez et al. 2016). This, in turn, suggests that both averages and variances will have an important impact on different performance related traits (e.g. Lardies et al. 2014; Vasseur et al. 2014; Bartheld et al. 2017). Nevertheless, we still do not know whether selection might also target traits as a function of those extremes.

In this context, populations inhabiting highly seasonal environments characterized also by daily extreme temperatures, provide a natural laboratory to evaluate the role of natural selection on the plasticity of critical thermal limits and preferences. We addressed such important gaps in our knowledge by measuring for the first time survival as a function of the plasticity of thermal critical temperatures (CT_Max_ and CT_Min_), preferred temperature (T_Pref_) and thermal sensitivity of metabolism (Q_10_; the magnitude of change in metabolic rate for a 10°C change in T_a_) after acclimation to 10**°**C and 20**°**C in the northernmost population of the four-eyed frog *Pleurodema thaul*. We tested four predictions regarding phenotypic selection and plasticity that built up from previous findings showing that acclimation to warmer temperatures produces an increase in the upper but not in the lower limits of the thermal performance curve (Ruiz-Aravena et al. 2014) (Fig. 1). First, the high seasonality should select for plasticity in TPC parameters and therefore, the plasticity itself should currently be under directional selection. Second, if daily high extreme temperatures were frequent, then we would expect positive directional selection on CT_max_ when warm as well as cold acclimated. Third, if daily low extremes were frequent, then we would expect negative directional selection on CT_min_ during the cooler periods of the year. Fourth, as energy inputs are limited, the energetic definition of fitness indicates that individuals with higher maintenance costs (i.e. resting metabolic rate) would have less energy available to allocate to growth, reproduction and/or performance. The main prediction of this principle is that natural selection should maximize the residual available energy, and therefore, higher maintenance costs would be associated with lower fitness if no compensations in other functions occur (Bacigalupe & Bozinovic 2002; Artacho & Nespolo 2009). Thus, our final prediction is that Q_10_ is not under directional selection.

**Figure 1.**
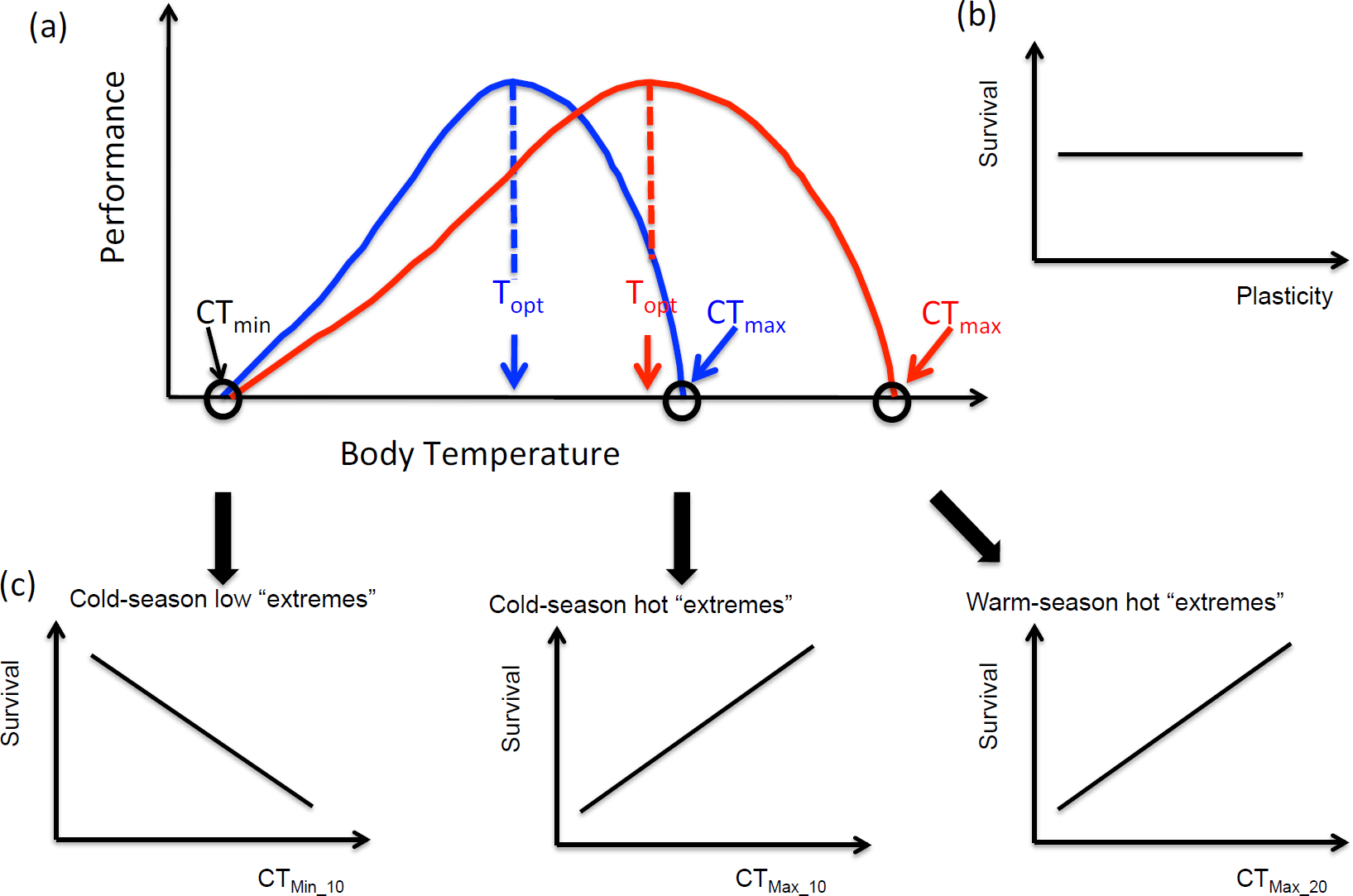
Graphical representation of the theory tested in this study. (a) Predictions built up from findings showing that acclimation to warmer temperatures produces an increase in the upper but not in the lower limits of the thermal performance curve (Ruiz-Aravena et al. 2014). (b) The high seasonality should selected for plasticity and therefore, plasticity of all traits should currently be under directional selection. (c) If daily low extremes are frequent, negative directional selection on CT_Min_ during the cooler periods of the year is expected (left panel). If daily high extreme temperatures are frequent, positive directional selection on CT_Max_ during the warmer periods (right panel) as well as the cooler periods of the year is expected (middle panel). We predict no directional selection on T_Pref_ and Q_10_ at both acclimation temperatures and on CT_Min_ when warm acclimated. Cold acclimation is indicated by a _10 subscript while warm acclimation is indicated by a _20 subscript.

## METHODS

### Study organism and laboratory maintenance

Eighty-three adults individuals of *P. thaul* were captured during September 2012 on two small ponds at Carrera Pinto (27°06’40.2” S, 69°53’44.3” W; 2,000 m.a.s.l.), a small oasis in the Atacama Desert that is known to be the northernmost population of the species (Correa et al. 2007). In both ponds, we performed an exhaustive search across microhabitats (below rocks, in the vegetation and in the water). All individuals were transported to the laboratory (Universidad Austral de Chile, Valdivia) within 2 – 3 days of capture. Following capture all animals were marked by toe clipping and maintained in the laboratory for one month at a temperature of 20° ± 2°C and with a photoperiod 12D:12L. Animals were housed (N = 5) in terraria (length × width × height: 40 × 20 × 20 cm) provided with a cover of moss and vegetation and a small recipient filled with water. Individuals were fed once a week with mealworms (*Tenebrio sp*. larvae) and Mazuri® gel diets.

### Acclimation and thermal traits

After one month at maintenance conditions, in a split cross design half the frogs were acclimated to either 10**°**C or 20**°**C for two weeks before measuring thermal traits. Frogs were randomly assigned to the first acclimation temperature using a coin. Next they were acclimated to the other temperature and again measured thermal traits. We chose these acclimation temperatures because they are close to the mean minimum temperatures during the breeding season (August – October, 10°C) and to the mean temperatures during the active period of the species (20°C) at Carrera Pinto (www.cr2.cl). None of the investigators were blinded to the group allocation during the experiments.

Critical temperatures were determined as the environmental temperature at which an individual was unable to achieve an upright position within 1 minute (Ruiz-Aravena et al. 2014). Each individual was placed in a small chamber inside a thermo-regulated bath (WRC-P8, Daihan, Korea) at 30°C (CT_Max_) or 5°C (CT_Min_) for 15 minutes, after which the bath temperature was increased (or decreased) at a rate of 0.8°C per minute (Rezende et al. 2011). Every minute or at every 1°C change, the chamber was turned upside down and we observed if the animal was able to return to the upright position. When an animal was unable to achieve an upright position within 1 minute it was allowed to recover at ambient temperature (CT_Min_) or for 30 minutes in a box with ice packs (CT_Max_). Body mass (a proxy of body size) was obtained before each trial using a Shimadzu TX323L electronic balance.

Preferred temperature (T_Pref_) was determined simultaneously for five individuals in five open-top terraria (length × width × height: 85 × 12 × 30 cm). Each terrarium had a thermal gradient between 10°C and 30°C produced by an infrared lamp overhead (250 W) on one end, and ice packs on the other. The organic gardening soil was moisturized at the beginning of each trial to prevent the desiccation of the frogs. Five individuals were placed at the centre of each one of the terraria and 45 minutes later we registered T_Pref_ as the dorsal body temperature (T_b_) using a UEi INF155 Scout1 infrared thermometer. Dorsal and cloacal T_b_ are highly associated (*r*_*P*_ = 0.99) (see Ruiz-Aravena et al. 2014 for details). Body mass was obtained before each trial using a Shimadzu TX323L electronic balance.

Standard metabolic rate, measured through oxygen consumption at 20°C and 30°C was measured continuously using an infrared O_2_ – CO_2_ analyzer (LI-COR LI6262, Lincoln, NV, USA). The analyzer was calibrated periodically against a precision gas mixture. Although there was almost no difference between calibrations, baseline measurements were performed before and after each recording. Flow rates of CO_2_ – free air was maintained at 100 ml min^−1^ ± 1% by a Sierra mass flow controller (Henderson, NV, USA). We used cylindrical metabolic chambers (60 ml) covered by metal paper. O_2_ consumption was recorded during 45 minutes per individual. Each record was automatically transformed by a macro program recorded in the ExpeData software (Sable Systems), to (1) transform the measure from % to mlO_2_ min^−1^, taking into account the flow rate and (2) to eliminate the first 5 min of recordings. For each individual, the metabolic sensitivity (Q_10_) was calculated as the ratio between metabolic rate measured at 30°C and metabolic rate measured at 20°C.

### Selection on thermal traits

After the experiments, all frogs were put back to 20°C for at least one month before releasing them. Marked frogs were released at Carrera Pinto in April 2013 and their survival was monitored on three separate recapture efforts (13^th^ October 2013, 13^th^ June and 9^th^ September 2014). As the desert surrounds these two small ponds dispersal was not a concern. The relationship between trait plasticity and survival was analyzed using a Cormack-Jolly-Seber (CJS) framework in Program MARK. An overall goodness of fit test was run using U-Care to ensure the data were consistent with the assumed structure of the CJS model and to obtain a value for the over dispersion parameter (c-hat). The time interval between capture occasions (as a fraction of 1 year and considering also the original capture event) was included in the analysis to accommodate the unequal intervals. The resulting resighting and survival estimates were therefore corrected to annual estimates. Survival and resighting parameters were obtained in a two-stage process. First, the best-fit resighting model was identified from three candidate models (constant, time dependent, and a linear trend). The fit of the three candidate resighting models was compared using survival modeled as both a constant and a time-dependent rate, to ensure that selection of the best-fit resighting model was not influenced by choice of survival model. Once the best-fit resighting model was identified (using AICc), it was then retained for all candidate models. A model selection and an information-theoretic approach (Burnham & Anderson 2003) was employed to contrast the adequacy of different working hypotheses (the candidate models) of selection on trait plasticity. The number of candidate models was kept to a minimum to minimize the likelihood of spurious results (Burnham & Anderson 2003; Lucaks et al. 2010). Body mass showed a positive relationship with CT_Max_20_ (*r*_*P*_ = 0.47) and with T_Pref_10_ (*r*_*P*_ = 0.24) but was not associated with any other trait (results not shown). Therefore, we tested only for a null model (i.e. neither trait under selection), a model with body mass and models with directional selection for each trait separately and also for correlational selection (interaction of trait combinations) in the same trait at both acclimation temperatures, which indicates plasticity. Body mass was included as a covariate in the case of CT_Max_20_ and T_Pref_10_ (Table 1). All analyses were performed in R version 3.1.3 employing package RMark (Laake 2013). No transformation was required to meet assumptions of statistical tests. Survival in relation to each covariate was obtained as the model averaged value across all candidate models weighted by individual model probability (Table 1).

**Table 1.** Candidate models ordered accordingly to their Akaike weights. Single term models represent directional selection (e.g. CT_Max_) and correlational selection represents plasticity (e.g. CT_Max_10_ * CT_Max_20_). CT_Min_ = minimum critical temperature; CT_Max_ = maximum critical temperature; T_Pref_ = preferred temperature; Q_10_ = thermal sensitivity of metabolism; MB = body mass. Cold acclimated is indicated by a _10 subscript while warm acclimated is indicated by a _20 subscript.

| Models | K | AICc | $\Delta AICc$ | $w_i$ |
| --- | --- | --- | --- | --- |
| 1 Null model | 2 | 130.17 | 0 | 0.220 |
| 2 $CT_{Min\_10}$ | 3 | 131.40 | 1.23 | 0.119 |
| 3 MB | 3 | 131.78 | 1.61 | 0.098 |
| 4 $T_{Pref\_20}$ | 3 | 132.08 | 1.90 | 0.085 |
| 5 $Q_{10\_10}$ | 3 | 132.18 | 2.01 | 0.081 |
| 6 $CT_{Min\_20}$ | 3 | 132.25 | 2.08 | 0.078 |
| 7 $CT_{Max\_10}$ | 3 | 132.26 | 2.08 | 0.078 |
| 8 $Q_{10\_20}$ | 3 | 132.26 | 2.09 | 0.077 |
| 9 $CT_{Min\_10} + CT_{Min\_20} + CT_{Min\_10} * CT_{Min\_20}$ | 5 | 133.38 | 3.21 | 0.044 |
| 10 $MB + CT_{Max\_20}$ | 4 | 133.44 | 3.27 | 0.043 |
| 11 $MB + T_{Pref\_10}$ | 4 | 133.82 | 3.64 | 0.036 |
| 12 $Q_{10\_10} + Q_{10\_20} + Q_{10\_10} * Q_{10\_20}$ | 5 | 134.17 | 4.00 | 0.030 |
| 13 $MB + T_{Pref\_10} + T_{Pref\_20} + T_{Pref\_10} * T_{Pref\_20}$ | 6 | 137.16 | 6.99 | 0.007 |
| 14 $MB + CT_{Max\_10} + CT_{Max\_20} + CT_{Max\_10} * CT_{Max\_20}$ | 6 | 137.62 | 7.45 | 0.005 |

$K$ = number of parameters.
AICc: AIC values corrected for small sample sizes.
$w_i$ : Akaike weights.

## RESULTS

All measured traits including critical thermal limits (CT_Max_, CT_Min_), thermal preference (T_Pref_) and sensitivity of metabolic rate to temperature (Q_10_) showed high variance among individuals (Fig. 2). In addition, for all traits some individuals shifted their thermal traits to higher values when acclimated to high temperatures, but other individuals showed the reverse response, that is their traits shifted to lower values after acclimation at higher temperatures (Fig. 3).

**Figure 2.**
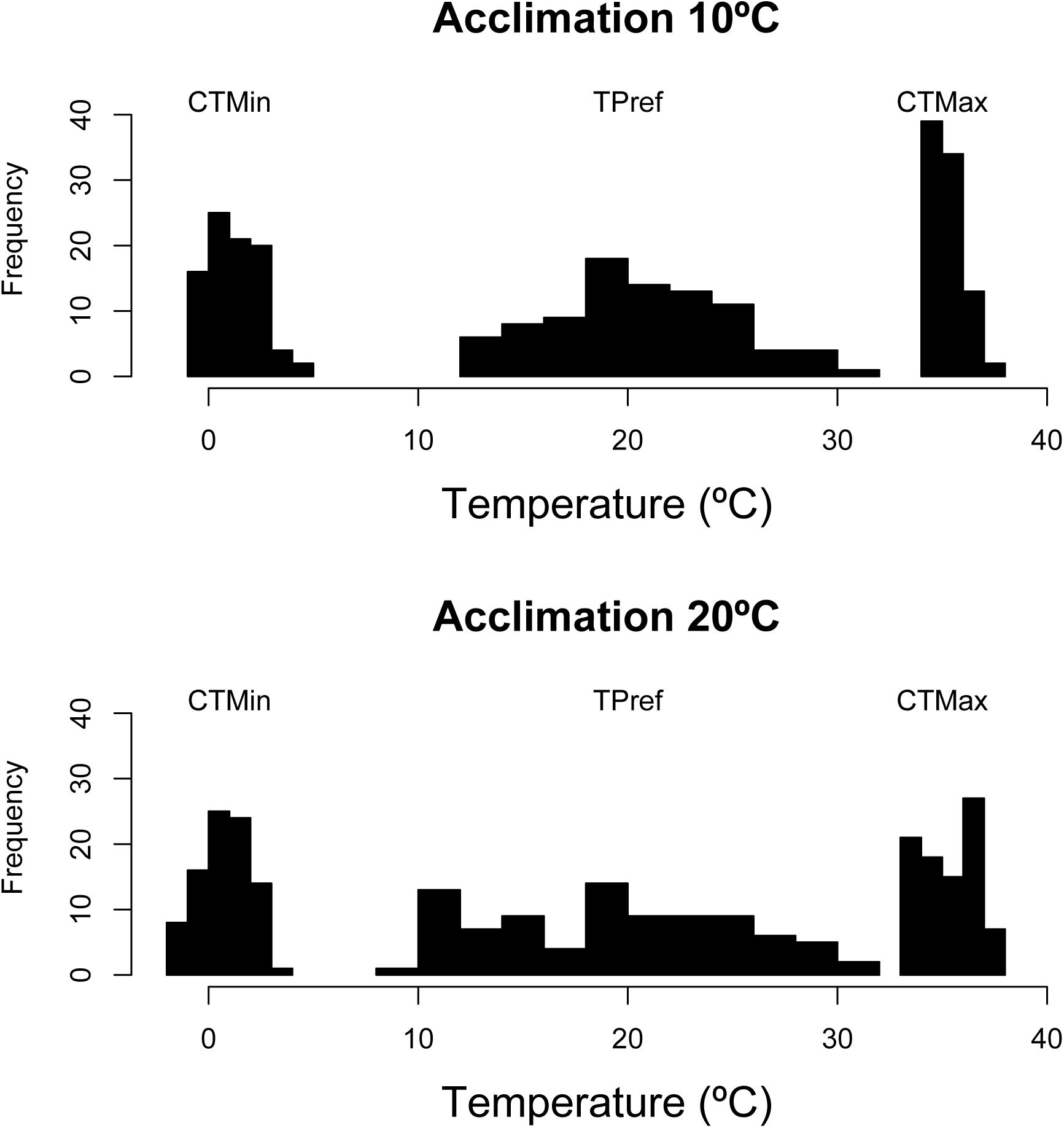
Frequency distribution of CT_Min_, T_Pref_ and CT_Max_ of the four-eyed frog when acclimated to 10°C and 20°C.

**Figure 3.**
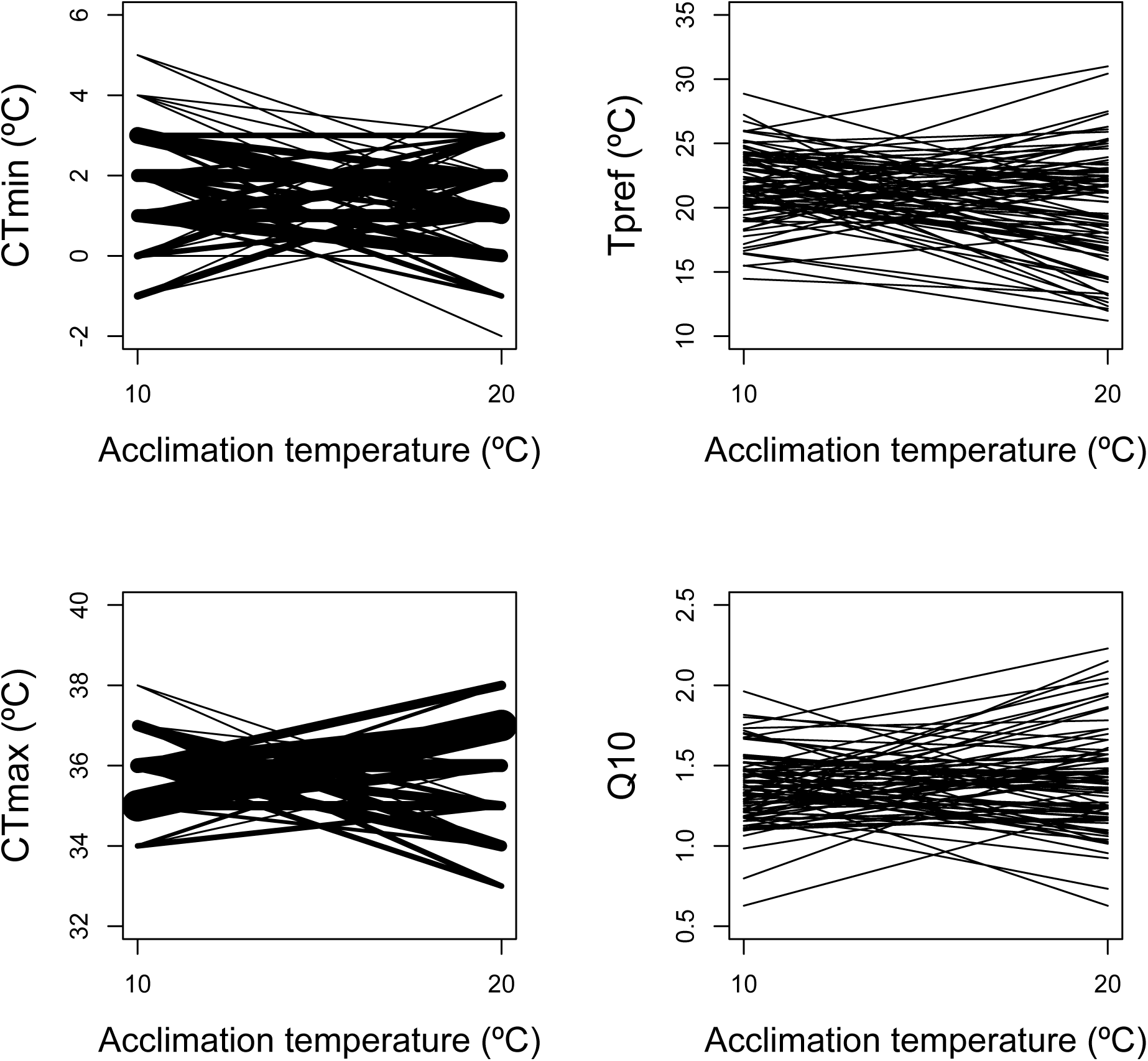
Individual plasticity in CT_Min_, T_Pref_, CT_Max_ and Q_10_ in response to 10 and 20°C acclimation treatments. Each line represents the individual value of the given trait at each acclimation temperature. For CT_Min_ and CT_Max_ the width of the line is directly proportional to the number of individuals that showed that specific response.

Only 5 out of 28 correlations between physiological traits were statistically significant, and these involved mostly critical thermal limits. In particular CT_Max_20_ was negatively correlated with CT_Min_10_ (*r*_*P*_ = −0.57) and CT_Max_10_ (*r*_*P*_ = −0.41) whilst it was positively correlated with Q_10_20_ (*r*_*P*_ = 0.26). Additionally, CT_Max_10_ was positively correlated with CT_Min_10_ (*r*_*P*_ = 0.31) and negatively correlated with CT_Min_20_ (*r*_*P*_ = −0.25).

The overall goodness of fit measure for the CJS model indicated a moderate level of over-dispersion (c-hat = 2.65, *P* = 0.103), however with only 3 recapture occasions it was not possible to identify an alternative starting model and the basic CJS model was adopted as the basis for subsequent model fitting, with unexplained over-dispersion controlled using the c-hat adjustment. A constant resighting rate was the best-fit model irrespective of whether survival was modeled as a constant or time dependent rate (Table 1). Consequently, the constant rate-resighting model was retained for subsequent modeling of survival. The model selection procedure indicated that from the 13 candidate models tested, there was not a single best-fit one (Table 1). In particular, the null model was the most supported (Akaike weight of 0.220), whilst models including only directional selection on single traits still had some support, with a cumulative Akaike weight of almost 60% (Table 1). Models including correlational selection (i.e. plasticity) showed rather weak empirical support (Table 1). Overall, survival decreased as values of most of the traits increased in both, warm and cold acclimated conditions (Table 2, Figure 4).

**Table 2.** Directional selection estimates from single terms models with their standard errors (SE) and 95% confidence intervals (95% CI). CT_Min_ = minimum critical temperature; CT_Max_ = maximum critical temperature; T_Pref_ = preferred temperature; Q_10_ = thermal sensitivity of metabolism; MB = body mass. Cold acclimation is indicated by a _10 subscript while warm acclimation is indicated by a _20 subscript.

| Trait | Estimate | SE | 95% CI |
| --- | --- | --- | --- |
| MB | 0.209 | 0.212 | -0.206 – 0.625 |
| $CT_{Min_{10}}$ | -0.248 | 0.187 | -0.616 – 0.119 |
| $CT_{Min_{20}}$ | -0.030 | 0.181 | -0.384 – 0.324 |
| $T_{Pref_{10}}$ | -0.025 | 0.059 | -0.140 – 0.090 |
| $T_{Pref_{20}}$ | -0.026 | 0.042 | -0.109 – 0.056 |
| $CT_{Max_{10}}$ | 0.026 | 0.257 | -0.477 – 0.530 |
| $CT_{Max_{20}}$ | -0.192 | 0.195 | -0.575 – 0.191 |
| $Q_{10_{10}}$ | -0.475 | 1.140 | -2.709 – 1.759 |
| $Q_{10_{20}}$ | -0.048 | 0.795 | -1.607 – 1.510 |

**Figure 4.**
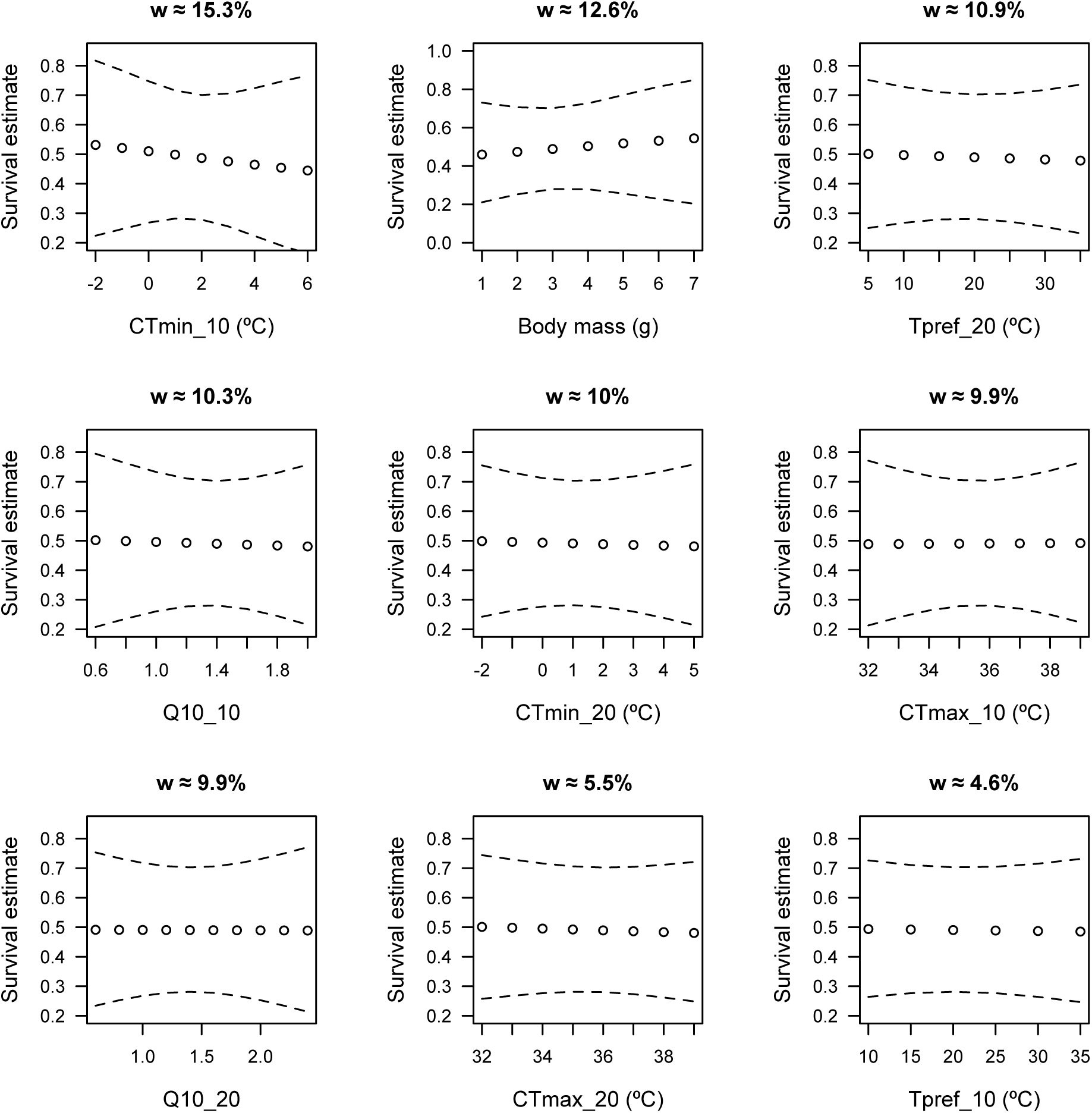
Directional survival selection on various thermal traits sorted by model probabilities. *w*: Akaike weights of the model.

## DISCUSSION

To understand how organisms adapt to highly fluctuating environments and whether they will be able to adaptively respond to current climate change, we need to evaluate whether selection in nature targets plasticity itself. Populations inhabiting highly seasonal environments that also experience daily extreme temperatures, provide excellent opportunities to test predictions of the fitness consequences of such thermal variation on the plasticity of critical thermal limits and preferences. Here, to the best of our knowledge for the first time, we studied natural selection on thermal acclimation capacity of performance (CT_Max_ and CT_Min_), metabolism (Q_10_) and behaviour (T_Pref_). Our results indicate that thermal acclimation in this population is not being targeted by directional selection, although there might be signals of selection on individual traits. In part, the relatively weak evidence for natural selection on this system might be a consequence of the small sample size we used (N = 83), the few recaptures we carried out (n = 3) and the relatively high value of c-hat in the analyses, which penalizes models on the basis of parameter number. This prevented us not only from evaluating more complex models (i.e. non linear selection) but also resulted in estimates of directional selection with rather large SEs and therefore with 95% confidence intervals that contained the zero in all cases.

Some theoretical models of thermal adaptation across heterogeneous environments (e.g., climate variability hypothesis, generalist-specialist models) suggest that temporal environmental heterogeneity selects for more broadly adapted individuals (Lynch and Gabriel, 1987; Gilchrist 1995), favoring increased plasticity particularly in thermal tolerance traits (Gunderson & Stillman 2015). Based on these models we predicted that the high seasonality should select for high plasticity in thermal traits and therefore, the plasticity itself should currently be under directional selection. Our prediction turned out to be incorrect as models including plasticity showed relatively weak support.

Frogs of *P. tahul* in the Atacama Desert, the northernmost population of this species, are exposed to large daily and seasonal oscillations in environmental temperatures. The ratio between daily and annual thermal ranges (O’Donnell & Ignizio 2012) experienced by this extreme population (0.65) is ca. 15% higher than that of a population 2,000 km south (0.52), which experiences narrower daily environmental temperatures at the center of the species’ distribution (Barria & Bacigalupe 2017). This means that the studied population experiences a daily variation that is almost 65% of its seasonal variation. This high daily variation, in combination with the fact that climate change is already changing the frequency and intensity of extreme temperatures (Rahmstorf & Coumou 2011; Wang & Dillon 2014; Vasquez et al. 2017), made us wonder whether selection in nature might also target thermal traits as a function of daily extremes. As the cooler end of the thermal performance curve did not change trough acclimation to warmer temperatures (Ruiz-Aravena et al. 2014) we expected negative directional selection on CT_min_ during the cooler but not the warmer periods of the year. Our results are in agreement with the trend specified by this prediction, as survival decreased as CT_min_ increased (i.e. less tolerance to cold) when cold-acclimated (albeit the estimate was non-significant), which was the second most supported model (Table 1).

Although acclimation produced an increase in the upper limits of the thermal performance curve in this population (Ruiz-Aravena et al. 2014), we expected positive directional selection on CT_max_ when warm as well as cold-acclimated if daily high extreme temperatures were frequent. Our results do not offer support for this prediction: there was a slight trend for survival to decrease as CT_max_ increased under warm as well as under cold-acclimated conditions. However, in both cases estimates were not statistically different from zero. Nevertheless, this might suggests that selection could be favouring individuals that avoid hot microhabitats, possibly by means of behavioural responses (Ruiz-Aravena et al. 2014). Indeed, behavioural thermoregulation has been proposed as one key factor that prevents an evolutionary response to selection to raising temperatures (Kearney et al. 2009; Huey et al. 2012; Buckley et al. 2015). The fact that CT_Max_20_ was negatively correlated with CT_Min_10_ indicates that individuals with higher cold tolerance might be the ones avoiding hot microhabitats, which opens very interesting questions for further research.

Regarding the sensitivity of metabolism to temperature (Q_10_) we expected that Q_10_ not to be under directional selection. Our results are in (partial) agreement with that expectation, as the rate at which survival changed with changes in Q_10_ was very small (Fig. 4, Table 2), although the models with Q_10_ still showed some support (Table 1). Finally, we also expected no directional selection on T_Pref_ as we have previously shown that acclimation to warmer temperatures produced an increase in this trait (Ruiz-Aravena et al. 2014). Nevertheless, we found a non-significant trend showing that survival decreased, although at a very low rate, as T_Pref_ increased, which might suggest that selection favours those individuals that are able to avoid hot microhabitats.

Our results indicate a positive trend of survival with body size (although the directional selection estimate was non-significant), something that has been previously reported in the literature (Aubin-Horth et al. 2005; Iida & Fujisaki 2007; Crosby & Latta 2013; Delaney & Warner 2017). This is somewhat unsurprising, given that body mass is known to be positively associated with several physiological traits that enhance performance (Castellano et al. 1999; Madsen & Shine 2000; Hurlbert et al. 2008; Shepherd et al. 2008; Luna et al. 2009) including plasticity itself (Whitman & Ananthakrishnan 2009). Our oasis population inhabits two highly isolated ponds where other anuran competitors have not been observed, but there might be a risk of predation by herons (*L.D.B. personal observation*), which could explain the positive selection for body size. Nevertheless, further experimental work is needed to evaluate this possibility.

It is important to mention that we here measured plasticity in only one life stage. Likely other ecological and physiological traits are also plastic in this species, and their responses to acclimation might differ, also among different life stages. However, we still consider our results show a signal and provide the first evidence that phenotypic plasticity is not an actual target of selection in nature, but that daily climate extremes might be selecting for higher tolerance. Nevertheless, further work including multiple traits and life stages and also in other populations, should help to strengthen the trends found here into further generic hypotheses to clarify the role of plasticity for the viability of ectotherm populations in nature.

## Acknowledgements

We thank Nadia Aubin-Horth, Wolf Blanckenhorn, Dries Bonte, Ray Huey and Michael Logan for highly valuable comments on a previous version on the manuscript. LDB wish to acknowledge the friendship and great support of Don Demetrio and Sra. Blanca at Carrera Pinto’s oasis.

## Data

data is available for download from the CSIRO Data Access Portal (https://data.csiro.au/dap/landingpage?pid=csiro:29733) Doi: 10.4225/08/5a9727318bd0f

## Author Contributions

L.D.B conceptualized the study, designed the experimental procedures and carried out the experiment with A.M.B., A.G.M., M.R.A. and J.D.G.E; M.T., B.S. and L.D. B. analyzed the data and L.D.B., B.S. and J.D.G. wrote the paper with input from A.M.B and M.R.A.

## Ethics

This study did not involve endangered or protected species and was carried out in strict accordance with the recommendations in the Guide for the Care and Use of Laboratory Animals of the Comisión Nacional de Investigación Científica y Tecnológica de Chile (CONICYT). All experiments were conducted according to current Chilean law. The protocol was approved by the Committee on the Ethics of Animal Experiments of the Universidad Austral de Chile.

